# Suppression of RIPK3 by EZH2 contributes to cigarette smoking-induced chemoresistance in lung cancer

**DOI:** 10.64898/2026.08.07.743587

**Authors:** Rong Liu, Aifang Zhang, Jeffrey Yang, Gutian Xiao, Dongshi Chen

## Abstract

Lung cancer remains the leading cause of cancer-related mortality worldwide, with cigarette smoking (CS) representing its primary risk factor. In addition to promoting tumorigenesis, chronic CS exposure contributes to chemotherapy resistance, although the underlying mechanisms remain poorly understood. Here, we established a long-term CS exposure model by repeatedly treating Lewis lung carcinoma (LLC) cells with cigarette smoke extract (CSE). After 4 months of exposure, CSE-treated cells exhibited enhanced proliferation, migration, and resistance to chemotherapy-induced cell death. Mechanistically, chronic CSE exposure suppressed receptor-interacting protein kinase 3 (RIPK3) expression by upregulating the epigenetic regulator enhancer of zeste homolog 2 (EZH2), which promoted repressive histone methylation at the RIPK3 promoter. Loss of RIPK3 impaired chemotherapy-induced cell death primarily by inhibiting ferroptosis rather than necroptosis. Importantly, genetic depletion or pharmacological inhibition of EZH2 restored RIPK3 expression and sensitized lung cancer cells to gemcitabine treatment both in vitro and in vivo. Furthermore, analysis of human lung cancer datasets revealed an inverse correlation between EZH2 and RIPK3 expression, with RIPK3 levels progressively decreasing with smoking history. Collectively, these findings identify the EZH2/RIPK3 axis as a critical mediator of smoking-associated chemoresistance and uncover a previously unrecognized role for RIPK3 in ferroptosis regulation. Targeting EZH2-mediated RIPK3 suppression may represent a promising therapeutic strategy to overcome chemoresistance in lung cancer patients with a history of smoking.

## Introduction

Lung cancer is the leading cause of cancer death in the United States and worldwide. As one of the most important risk factors, long-term cigarette smoking (CS) increases tumor progression and cancer therapy resistance[1, 2]. CS exposure leads to mutation or epigenetic changes of tumor suppressors, enabling cancer cells to evade cell death-related pathways, which is an important mechanism in lung tumor progression and resistance to cancer therapy[3, 4].

Previous studies mostly focused on the mechanism of apoptosis suppression by CS in lung cancer tumorigenesis and chemoresistance[4]. However, the tremendous efforts on enhancing apoptosis-related mechanisms have only moderately improved lung cancer chemotherapy[5]. Increasing evidence suggests that alternative forms of programmed cell death, including necroptosis and ferroptosis, can suppress tumor progression and may provide therapeutic opportunities to overcome apoptosis resistance[6–9]. In addition to their cytotoxic effects, these non-apoptotic cell death pathways can stimulate antitumor immune responses, further enhancing their therapeutic potential[5, 10, 11]. However, it remains unclear if these non-apoptotic cell death mechanisms are involved in the long-term CS exposure-induced chemoresistance. In this study, we demonstrate that chronic CS exposure promotes lung cancer progression and chemoresistance by suppressing cell death mediated by receptor-interacting protein kinase 3 (RIPK3), the core mediator of necroptosis that is increasingly recognized to have critical, cross-pathway interactions with ferroptosis (6,14,18). We identify the key epigenetic regulator, enhancer of zeste homolog 2 (EZH2), as a crucial repressor of RIPK3 expression through promoting promoter-associated histone methylation. Importantly, restoration of RIPK3 expression via EZH2 knockdown or pharmacological inhibition re-sensitizes lung cancer cells to chemotherapeutic agents both *in vitro* and *in vivo*. Notably, our findings suggest that RIPK3 suppression primarily impairs chemotherapy-induced ferroptosis rather than necroptosis, further supporting the alternative role for RIPK3 in regulating other cell death pathways. Together, these findings uncover a previously unrecognized mechanism linking long-term CS exposure to chemoresistance through epigenetic silencing of RIPK3. Targeting the EZH2–RIPK3 axis to restore cell death sensitivity may represent a promising therapeutic strategy for lung cancer patients with a history of smoking.

## Materials and methods

### Cell culture and reagents

The Lewis lung carcinoma (LLC) cells (ATCC) were cultured in DMEM (Lonza, Pittsburgh, PA) containing 10 % FBS and 1% Penicillin/Streptomycin. The human lung cancer cell lines, A549, H1299, H460, H1650, HCC827 (ATCC), and H358, H1838 [12] were cultured in RPMI (Lonza, Pittsburgh, PA) supplemented with 10% FBS and 1% Penicillin/Streptomycin. All of the cell lines were routinely tested for mycoplasma contamination using PCR-based assays. All experiments were conducted using mycoplasma-free cells.

Cisplatin, gemcitabine (Gem), etoposide (Ept), and 5-fluorouracil (5-Fu), glutathione (GSH, Sigma-Aldrich) were dissolved in saline to prepare the stock solution. The Ezh2 inhibitor, GSK126, RIPK1 inhibitor, necrostatin-1 (Nec-1), MLKL inhibitor, Necrosulfonamide (NSA), caspases inhibitor, z-VAD-FMK (z-VAD), Ferroptosis inhibitor, Liproxstatin-1 (Lip1) were purchased from MedChemExpress (MCE, Monmouth Junction, NJ) and dissolved in DMSO to prepare stock.

### Generation of long-term CS exposure cells

The CSE was prepared using Kentucky 3R4F research-reference filtered cigarettes (The Tobacco Research Institute, University of Kentucky, Lexington, KY) as previously described[13]. Briefly, smoke from one cigarette was bubbled into 10 mL of cell culture medium to generate 100% CSE. LLC cells were exposed to 4% CSE for 8 h daily, followed by replacement with fresh culture medium for the remaining 16 h. Cells were passaged upon reaching >90% confluence. Chronic CSE exposure was maintained for periods ranging from 1 to 6 months.

### Cell viability and cell death assays

Cell viability was assessed using MTT assays (Promega, Madison, WI) and crystal violet staining. For MTT assays, cells were seeded in 96-well plates (1 × 10⁴ cells/well), treated with chemotherapeutic agents for 24 h, and analyzed using a Varioskan LUX microplate reader (Thermo Fisher Scientific).

For crystal violet staining, cells were seeded in 12-well plates (40–50% confluence), treated for 24 h, fixed with 3.7% paraformaldehyde, and stained with 0.05% crystal violet.

Cell death was further evaluated using Annexin-V-FITC/7-AAD staining (Invitrogen, Carlsbad, CA) followed by flow cytometry (BD FACSVerse).

### Edu staining, Reactive oxygen species (ROS), and transwell migration assays

Cell proliferation was assessed using the EdU Cell Proliferation Assay Kit (APExBIO) according to the manufacturer’s instructions. Briefly, cells were seeded in 12-well plates and cultured under the indicated conditions. Cells were incubated with 10 μM EdU for 6 h at 37°C, fixed with 4% paraformaldehyde, permeabilized with 0.5% Triton X-100, and stained using the Click-iT reaction cocktail. Nuclei were counterstained with DAPI. Images were acquired using a fluorescence microscope.

Intracellular ROS levels were analyzed using 2′, 7′-dichlorofluorescein diacetate (H_2_DCFDA) ROS assay kit (Tocris Bioscience, Minneapolis, MN) according to the manufacturer’s instructions. Briefly, cells were cultured at a density of 1 × 10^5^/well in 12-well plates or 1×10^4^ in the 96-well plates. After 24 h, cells were incubated with H_2_DCFDA at 37 °C for 30 min. Afterward, ROS levels in cells in 12-well plates were measured by flow cytometry (BD FACSverse) following trypsinization.

Cell migration was evaluated using Transwell chambers with 8-μm pore polycarbonate membranes (Corning). Briefly, 2.5 × 10^4^ cells suspended in serum-free medium were seeded into the upper chamber, while the lower chamber was filled with complete medium containing 10% fetal bovine serum as a chemoattractant. After incubation for 24 h at 37°C, non-migrated cells on the upper surface of the membrane were removed using a cotton swab. Cells that migrated to the lower surface were fixed with 4% paraformaldehyde, stained with 0.1% crystal violet. Images were acquired using a light microscope.

### Western blotting

Western blotting was conducted as previously described[14]. The following antibodies were used: RIPK3 (ab56164, Abcam, Waltham, MA) and (#2283, ProSci, Poway, CA), Gpx4 (#52455), MLKL (#14993), Ki-67 (#12202), Ezh2 (#5246), Snail (#3879), mouse p-MLKL (S345, #37333), cleaved caspase 3 (C Casp3, #9661), GAPDH (#2118, Cell Signaling, Danvers, MA), ACSL4 (sc-36523, Santa Cruz Biotechnology).

### RNA isolation and qRT-PCR

Total RNA was extracted using the Quick-RNA MiniPrep Kit (Zymo Research, Irvine, CA). cDNA was synthesized from 1 μg RNA using SuperScript II reverse transcriptase (Invitrogen). Quantitative PCR was performed for Ripk3, Ezh2, and GAPDH using primers as follows: mouse Ripk3 Forward, GAAGACACGGCACTCCTTGGTA; mouse Ripk3 Reverse: CTTGAGGCAGTAGTTCTTGGTGG; mouse Ezh2 Forward, CATACGCTCTTCTGTCGACGATG; mouse Ezh2 Reverse: ACACTGTGGTCCACAAGGCTTG; mouse Gapdh Forward, CGACTTCAACAGCAACTCCCACTCTTCC; mouse Gapdh Reverse, TGGGTGGTCCAGGGTTTCTTACTCCTT.

### Plasmid transfection and lentivirus infection

Plasmid transfections were performed using Lipofectamine 2000 (Thermo Fisher Scientific) according to the manufacturer’s protocol. The pcDNA3.1-HA-RIPK3 (#78804) for human and pcDNA3.1-HA-Ripk3 (#77805) for mouse RIPK3 plasmids were purchased from Addgene.

For the lentivirus infection, lentiviral particles were generated by co-transfecting 293T cells with the lentiviral vectors pMD2.G (VSVG, #12259), pMDLg/pRRE (#12251), and pRSV-REV (#12253) (Addgene)[14]. The pLKO.1 shRNA targeting mouse Ezh2 (TRCN0000304506) was purchased from Sigma-Aldrich. The pHAGE-EZH2 (Addgene #116738) was a gift from Dr. Jessie Huang at the University of Southern California. Stable cell lines were selected using puromycin (2 μg/mL) and validated by Western blotting.

### Bioinformatics analysis

To study the effect of cigarette smoking on gene expression changes in lung cancer patients, the gene expression profile dataset was downloaded from The Cancer Genome Atlas (TCGA) lung cancer database. The lung cancer patients were grouped by their smoking history (never smoker, n=93 vs current smoker, n=252). The data were analyzed using R (Version 4.1, http://www.bioconductor.org) with the edgeR package (SCR_012802). Fold change (FC) of gene expression was calculated with a cut-off of abs (log2FC) > (mean(abs(logFC))) and *P* value < 0.05.

The RIPK3, EZH2 expression in TCGA databases was analyzed by using the UCSC Cancer Genomics Browser (https://xenabrowser.net/)[15]. The correlation between RIPK3 and EZH2 expression and the overall survival of lung cancer patients was analyzed using the Kaplan-Meier Plotter (https://kmplot.com/analysis).

### Mice *in vivo* experiment

All animal studies were conducted with approval of the Institutional Animal Care and Use Committee Statement (IACUC) at the University of Southern California. Wild-type female C57BL/6J mice (JAX:000664) at 6∼8 weeks were subcutaneously injected with LLC cells (1 × 10⁶). After 7 days, mice were treated with gemcitabine (25 mg/kg, three times per week) and/or an EZH2 inhibitor (GSK126, 10 mg/kg, twice per week). Tumor volumes were measured using calipers and calculated as 0.5 × length × width². The maximal tumor volume did not exceed 4 cm³ under the permission of the institute IACUC. Tumor tissues were processed for flow cytometry or fixed in 10% formalin for immunofluorescence staining. Paraffin-embedded sections were stained with antibodies against GPX4 (#52455, Cell Signaling) and Cd8 (14-0081-85, Thermofisher), followed by fluorescent secondary antibodies and imaging using a Leica TCS SP8 confocal microscope.

Single-cell suspensions from tumors or lungs were prepared by enzymatic digestion (Dispase, collagenase D, and DNase I), followed by filtration and red blood cell lysis. Cells were stained with antibodies against CD45 (FITC, #157214), CD3 (Pe-Cy5, #300410), CD4 (PE-Cy7, #10042), and CD8 (Pacific Blue, #100725) (Biolegend) for surface markers. Data were acquired on a BD LSRFortessa and analyzed using FlowJo software (SCR_008520).

### Statistical Analysis

Statistical analyses were conducted using GraphPad Prism 8 software (GraphPad Software, Inc., La Jolla, CA). *P-*values were calculated using the student’s *t*-test and were considered significant if *P* <0.05. The means ± one standard error of the mean (SD) were displayed in the figures.

## Results

### Long-term CS exposure promotes lung cancer progression and chemoresistance

To investigate the impact of CS exposure on lung tumor progression, we exposed the LLC mouse lung cancer cells to long-term CSE (4% CSE, 8 h/day for 4 months, LLC_4M). Compared with parental LLC cells, LLC_4M exhibited significantly higher viability upon acute CSE (10% CSE) treatment for 16 h (Fig. S1A, B), suggesting that LLC cells developed resistance after long-term CSE exposure. We next evaluated the proliferative and migratory capacities of these cells using MTT, colony formation, Edu staining, and transwell migration assays. LLC_4M cells displayed enhanced proliferation and migration compared with their parental cells (LLC_P) (Fig. 1A–D). Consistently, the expression of proliferation marker Ki-67 and migration marker Snail was elevated in LLC_4M cells (Fig. 1E). To determine whether CS exposure confers resistance to chemotherapy, we treated both LLC_P and LLC_4M with multiple chemotherapeutic agents, including gemcitabine, cisplatin, etoposide, and 5-Fu. LLC_4M cells showed reduced sensitivity to all tested agents (Fig. 1F, G). To validate these findings *in vivo*, LLC_P and LLC_4M cells were implanted subcutaneously into C57BL/6J mice, followed by gemcitabine treatment, a first-line chemotherapy drug for lung cancer[16]. Tumors derived from LLC_4M cells grew more rapidly than those from LLC_P cells (Fig. 1H). While gemcitabine effectively suppressed LLC_P tumor growth, its inhibitory effect was markedly attenuated in LLC_4M tumors (Fig. 1H). Collectively, these results indicate that long-term CS exposure promotes lung cancer cell survival, enhances proliferative and migratory capacities, and confers resistance to chemotherapy.

**Figure 1.**
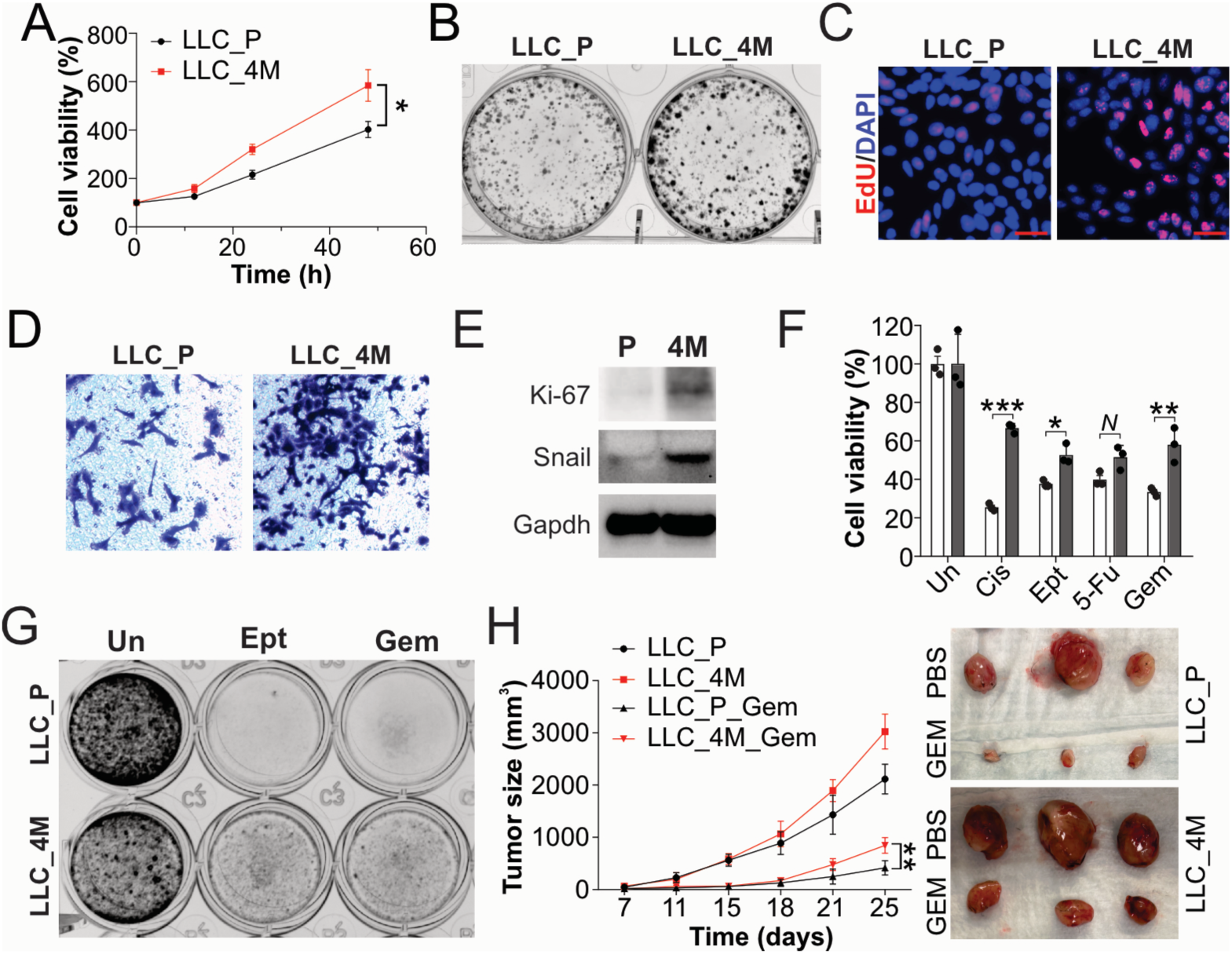
Long-term CSE exposure leads to chemoresistance. (A-E) LLC cells were exposed to 4% CSE for 8 hours/day over 4 months. (A) The growth of LLC parental (LLC_P) and 4-month CSE exposure (LLC_4M) cells was measured by MTT assay at different time points. (B) The colony formation of LLC_P and LLC_4M cells. (C) The EdU staining of LLC_P and LLC_4M cells. Scale bar, 25 μm. (D) The cell migration of LLC_P and LLC_4M cells was analyzed by the transwell migration assay. (E) The western blot of the indicated proteins in LLC_P and LLC_4M cells. (F, G) The LLC_P and LLC_4M cells were treated with cisplatin (Cis, 40 μM), etoposide (50 μM), 5-Fu (50 μg/ml), Gemcitabine (Gem, 200 ng/ml) for 24 h. Cell viability was analyzed using the MTT assay (F) and crystal violet staining (G). (H) WT female C56B/6J mice were subcutaneously injected with LLC_P and LLC_4M cells (1X10^6^) (n=5 for each group). 7 days later, the mice were treated with Gem (25 mg/kg, 3 times/week). The tumor size was measured on the indicated days. *Left*: growth curves were plotted; *right*: representative images of the tumors in each group. The experiments for (A-G) were repeated 3 times. Values in bar graphs are presented as means ± SD. N, p>0.05; *, p<0.05; **, p<0.01, ***, p<0.001.

### Long-term CS-exposed lung tumor cells have low RIPK3 expression

Since CS long-term exposure leads to resistance to CSE exposure and chemotherapy drugs, we hypothesized that this is due to dysregulation of the cell death signaling pathway. The flow cytometry data showed that Annexin-V+/7AAD- cells and 7AAD+ cells were suppressed in LLC_4M cells upon CSE treatment compared with their parental cells (Fig. 2A). We then systematically analyzed changes in cell death-related genes, including apoptosis, necroptosis, ferroptosis, pyroptosis, and autophagy, in lung cancer patients who were never smokers (n=93) and current smokers (n=252) using TCGA data. Among the 39 significantly differentially expressed genes, the downregulation of RIPK3 caught our interest (Fig. 2B). RIPK3 expression was inversely correlated with smoking history in lung cancer patients, showing progressive suppression in ever smokers, with the lowest levels observed in current smokers (Fig. 2C). We further confirmed the downregulation of RIPK3 in the LLC cell with different periods of CSE exposure at the protein and mRNA levels (Fig. 2D, E). The TCGA database analysis showed that RIPK3 is suppressed at the mRNA level in lung tumors when compared with normal solid tissue (Fig. S2A). Actually, the expression level of RIPK3 is also lower in lung cancer cells than in normal human bronchial epithelial cells (Fig. S2B). Using the Kaplan-Meier plotter (https://kmplot.com/analysis/), we found that lung cancer patients with higher RIPK3 expression have better clinical survival (Fig. 2F). These results collectively suggest that long-term CS exposure suppresses RIPK3 expression, which might promote lung tumor development and chemoresistance.

**Fig. 2.**
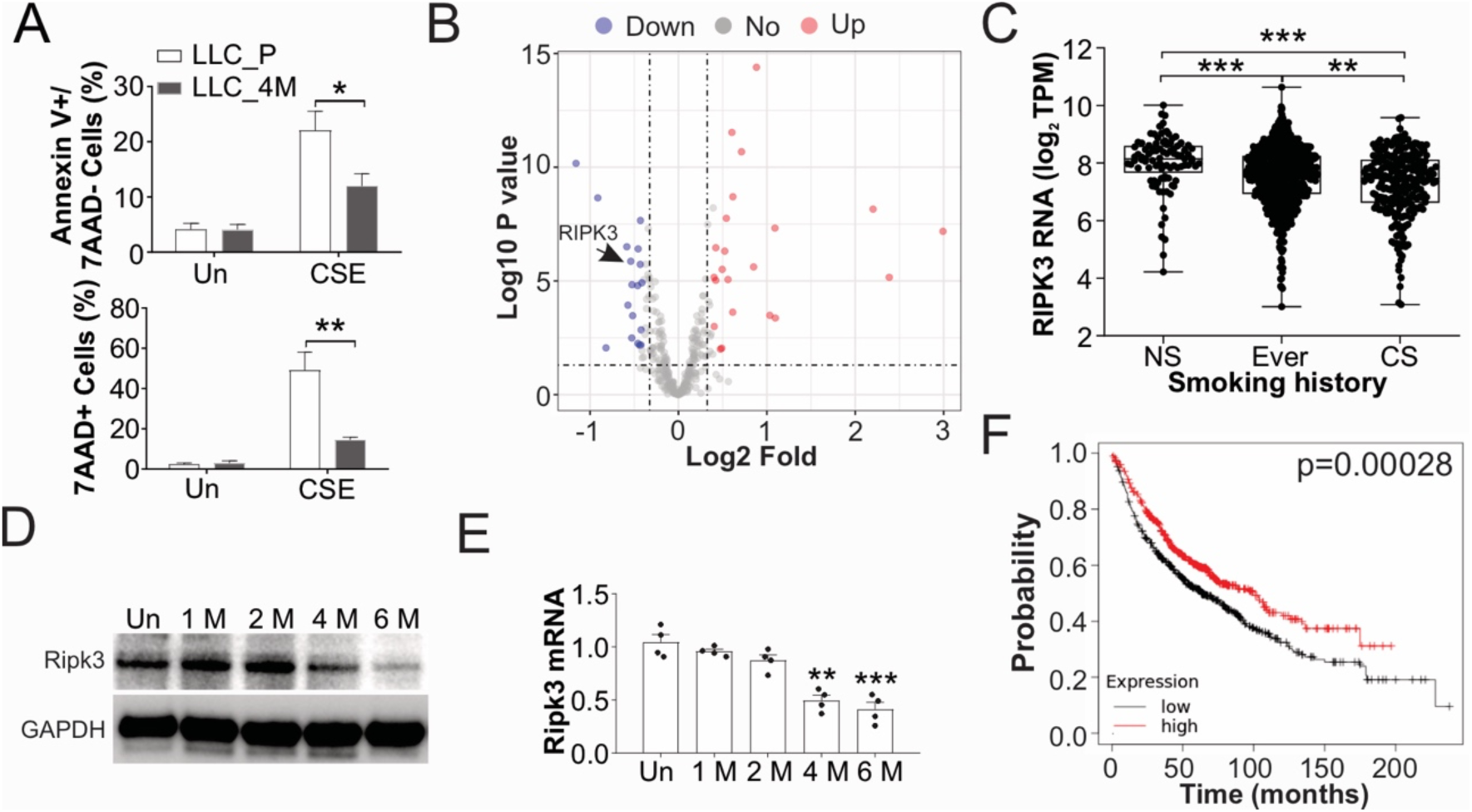
Long-term CS exposure suppresses the expression of RIPK3. (A) The LLC_P and LLC_4M cells were treated with 10 % CSE for 24 h. The cells were stained with PI and Annexin-V-FITC/7-AAD and analyzed by flow cytometry. The Annexin V+/7AAD- cells and 7AAD+ cells were quantified. (B) Differential expression of genes related to cell death between never smokers and current smokers in lung cancer patients, based on the TCGA database. (C) The mRNA level of RIPK3 in lung cancer patients of never smokers (NS, n=93), ever smokers (n=636), and current smokers (CS, n=252), based on the TCGA database. (D, E) LLC cells were exposed to 4% CSE for 8 hours/day over 1, 2, 4, and 6 months. The expression of Ripk3 in protein (D) and mRNA level (E) was analyzed. (F) Kaplan-Meier curves comparing overall survival (OS) in patients with lung tumors with high or low RIPK3 expression (https://kmplot.com/analysis/). The experiments for (A, D, E) were repeated 3 times. Values in bar graphs are presented as means ± SD. N, p>0.05; *, p<0.05; **, p<0.01, ***, p<0.001.

### Low expression of RIPK3 contributes to CS-induced lung tumor growth and chemoresistance

To test whether RIPK3 is involved in CS-induced tumorigenesis and chemoresistance, we re-expressed RIPK3 in LLC_4M cells by transfection with the pcDNA3.1-HA-RIPK3 plasmid. Recovery of Ripk3 expression suppressed the LLC_4M cells proliferation and migration (Fig. 3A), as well as suppressed the induction of Ki-67 and Snail (Fig. 3B). Moreover, Ripk3 overexpression re-sensitized the LLC_4M cells to gemcitabine and etoposide-induced cell death (Fig. 3C, D). Consistently, recovery of RIPK3 expression in human H460 lung tumor cells, which has lower RIPK3 expression (Fig. S2B), also enhances the killing effect of gemcitabine and etoposide (Fig. 3E, F). These data indicate that the suppression of RIPK3 expression by long-term CS exposure contributes to lung tumor cell growth and chemoresistance.

**Fig. 3.**
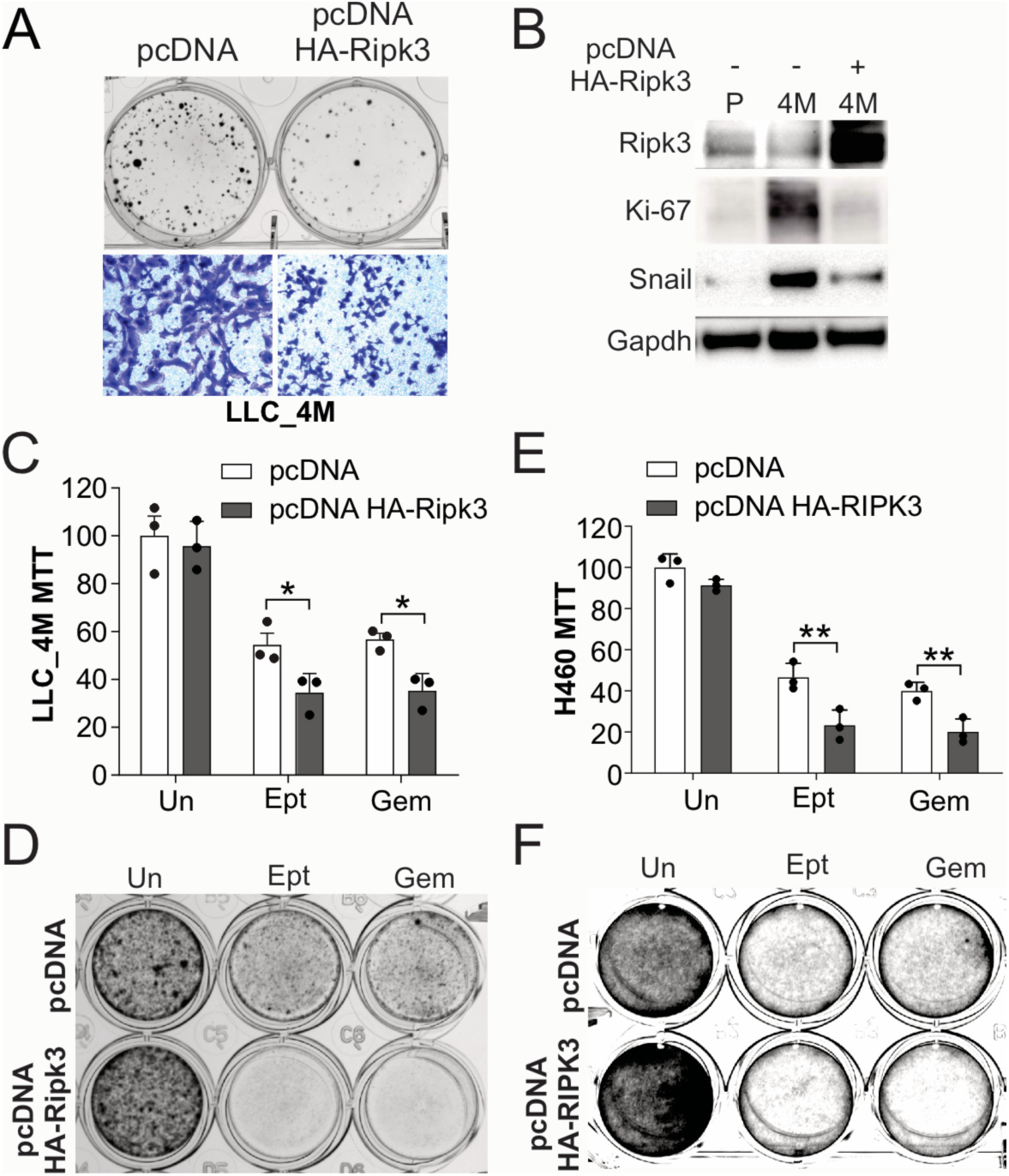
Enhanced expression of RIPK3 suppresses cell progression and chemoresistance. (A) The LLC_4M cells were transfected with pcDNA3.1 or pcDNA3.1 HA-Ripk3. *Upper*: crystal violet staining of colony formation at 14 days; *lower*, cell migration was analyzed by transwell assay. (B) The LLC_P and LLC_4M cells were transfected with pcDNA3.1 or pcDNA3.1 HA-Ripk3 for 24h. The expression of indicated proteins was analyzed by Western blot. (C, D) The LLC_4M cells transfected with pcDNA3.1 or pcDNA3.1 HA-Ripk3 were treated with Ept (50 μM) or Gem (200 nM) for 24h. Cell viability was assessed by the MTT assay (C) and crystal violet staining (D). (E, F) The H460 cells transfected with pcDNA3.1 or pcDNA3.1 HA-RIPK3 were treated with Ept (50 μM) or Gem (20 μM) for 24h. Cell viability was assessed by the MTT assay (E) and crystal violet staining (F). Each experiment was repeated 3 times. Values in bar graphs are presented as means ± SD. *, p<0.05; **, p<0.01.

### RIPK3 expression contributes to gemcitabine-induced ferroptosis

Since RIPK3 is an essential mediator of necroptosis (6, 14), we further investigated whether long-term CS exposure suppresses gemcitabine-based chemotherapy by inhibiting RIPK3-mediated necroptosis. Flow cytometry showed that LLC_4M cells have fewer 7AAD+ cells after gemcitabine treatment than parental cells (Fig. 4A). However, we did not observe the induction of p-MLKL, the final executor of necroptosis[17], upon gemcitabine treatment (Fig. 4B). Inhibition of MLKL by NSA or RIPK1 by Nec-1 did not compromise the gemcitabine-induced cell death in the H460 cells either (Fig. S3A). In addition to necroptosis, RIPK3 has been reported to mediate ferroptosis[18], which can also lead to 7AAD+ cells. We then tested ferroptosis markers, including ACSL4 and GPX4. While we don’t see a change in ASCL4, we found that GPX4 was suppressed by gemcitabine treatment in LLC_P cells (Fig. 4B), which was compromised in LLC_4M cells (Fig. 4B). And the supplement of ferroptosis inhibitor, Lip1, suppressed gemcitabine-induced cell death (Fig. S3A). Enhanced expression of RIPK3 in LLC_4M cells increased 7AAD+ cells, and suppression of Gpx4 upon gemcitabine treatment (Fig. 4C, D). While LLC_4M cells have less cleaved caspase-3 expression upon gemcitabine treatment (Fig. 4B), and supplementation of the apoptosis inhibitor, z-VAD, suppressed gemcitabine-induced cell death (Fig. S3A), the enhanced RIPK3 expression did not recover the cleavage of caspase-3 (Fig. 4D), suggesting that suppression of RIPK3 in the LLC_4M cells did not contribute to resistance to apoptosis. Since Ripk3 has been reported to induce ROS (19), which contributes to ferroptosis, we further investigated changes in ROS in LLC_P and LLC_4M cells. Here, we found that gemcitabine induced an increase in ROS in LLC_P cells, which was compromised in LLC_4M cells (Fig. 4E). Enhanced RIPK3 expression restored ROS levels in LLC_4M cells upon GEM treatment (Fig. 4E). Supplementation with the ROS scavenger, GSH, compromised the GEM-induced ROS (Fig. S3B) and cell death (Fig. 4F, S3A) in LLC_4M with Ripk3 overexpression. To test Gem-induced ferroptosis *in vivo*, we stained for Gpx4 in LLC_P and LLC_4M tumors. Consistently, we found that gemcitabine treatment suppressed Gpx4 in LLC_P tumors, but not in LLC_4M tumors (Fig. 4G). Collectively, our data suggested that long-term CS exposure suppressed RIPK3 and compromised chemotherapy drug-induced ferroptosis, thereby conferring chemoresistance.

**Fig. 4.**
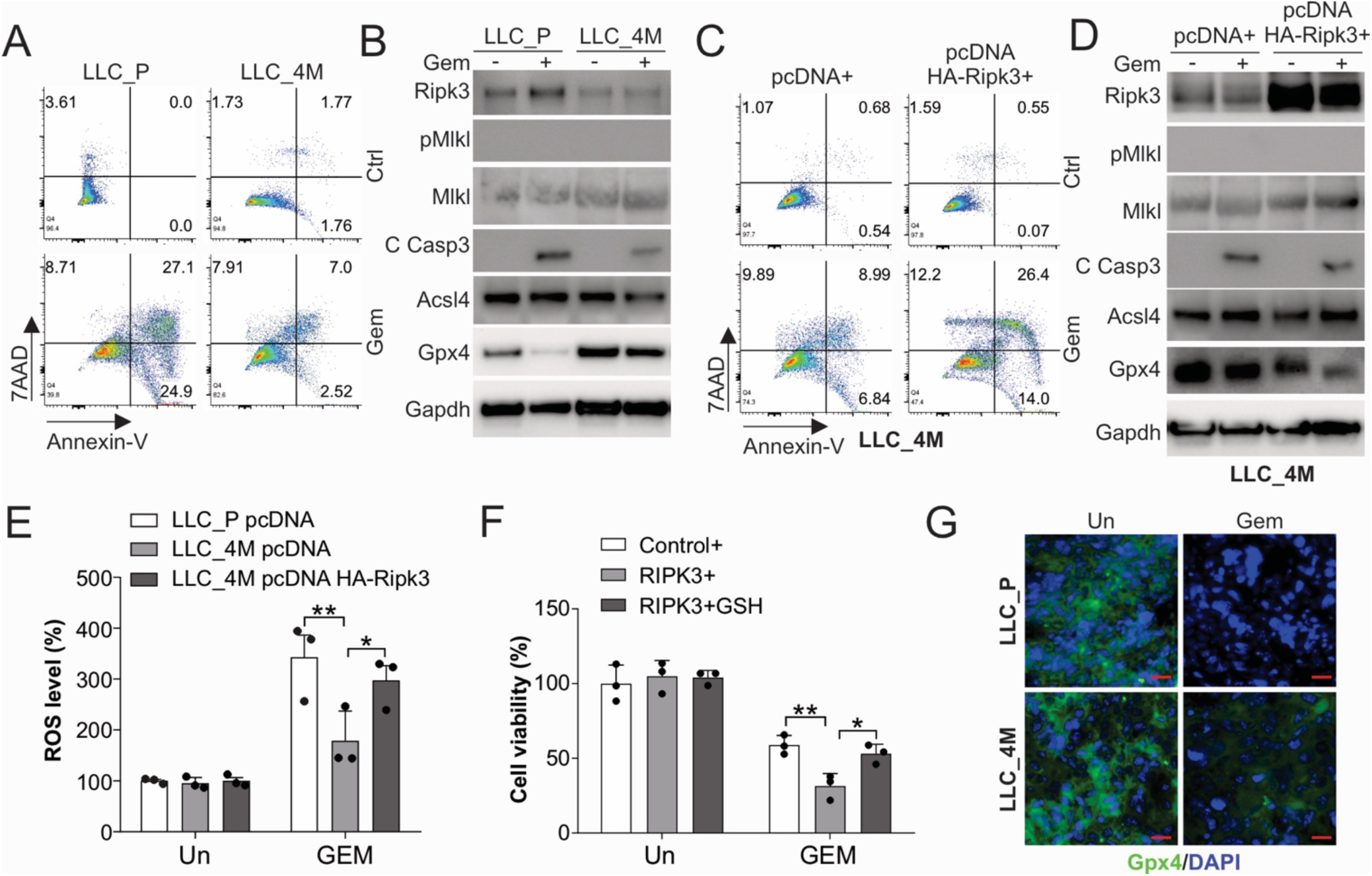
Enhanced RIPK3 sensitized the resistant lung tumor cells to chemotherapy-induced ferroptosis. (A, B) The LLC_P and LLC_4M cells were treated with 200 ng/ml Gem for 24h. (A) The cell death was analyzed by flow cytometry following Annexin-V-FITC/7AAD staining. (B) The expression of the indicated proteins was analyzed by western blot. (C, D) The LLC_4M cells transfected with pcDNA3.1 or pcDNA3.1 HA-Ripk3 were treated with 200 ng/ml Gem for 24h. (C) The cell death was analyzed by flow cytometry following Annexin-V-FITC/7-AAD staining. (D) The expression of the indicated proteins was analyzed by western blot. (E) The LLC_P and LLC_4M cells transfected with pcDNA3.1 or pcDNA3.1 HA-Ripk3 were treated with 200 ng/ml Gem for 24h. The ROS level in cells was measured using the H2DCFDA assay. (F) The LLC_4M cells transfected with pcDNA3.1 or pcDNA3.1 HA-Ripk3 were treated with 200 ng/ml Gem in combination with GSH for 24h. Cell viability was assessed by the MTT assay. (G) The Gpx4 staining in the LLC_P and LLC_4M tumors with or without Gem treatment. Scale bar, 25 μm. Each experiment was repeated 3 times. Values in bar graphs are presented as means ± SD. *, p<0.05; **, p<0.01.

### EZH2 suppresses RIPK3 expression in lung cancer cells upon long-term cs exposure

As RIPK3 has been reported to be promoter-associated epigenetically silenced [19], and CS exposure is a major source of DNA epigenetic modifications[20], we hypothesized that long-term CS exposure suppresses RIPK3 expression through epigenetic modifications. By analyzing the expression of epigenetic regulator genes [21] in lung tumors from never smokers (n = 93) and current smokers (n = 252), we observed that most epigenetic regulators were upregulated in current smokers (Fig. S4A). Among the significantly altered genes, EZH2 showed one of the greatest increases in expression and displayed the strongest negative correlation with RIPK3 expression (Fig. 5A, B, S4A), making it a compelling candidate for mediating RIPK3 repression in smoking-associated lung cancer. Opposite to RIPK3, the EZH2 expression is positively correlated with the smoking history in lung cancer patients (Fig. 5C). And higher EZH2 expression is correlated with poorer survival of lung cancer patients (Fig. 5D). We further confirmed that induction of EZH2 in the LLC cells with 4 months or 6 months CSE exposure in the protein and mRNA level (Fig. 5E). Since EZH2 catalyzes trimethylation of lysine 27 on histone H3 (H3K27me3), we analyzed several public databases (GSE75903[22], GSE249645, GSE164247[23], GSE307329[24], GSE291333[25]) and found that the H3K27me3 was enriched at RIPK3 promoter in lung cancer cell lines, including A549, H1299, and H1963 cells, but showed lower enrichment in normal lung epithelial cells, including alveolar type I (AT1) and II (AT2) cells, the cells of origin of lung adenocarcinoma (Fig. 5F). To test the effect of EZH2 expression on RIPK3 levels, we knocked down Ezh2 in LLC_4M cells using shRNA. We found that the absence of Ezh2 restored Ripk3 expression at the protein and mRNA levels (Fig. 5G, S4B). Knockdown of Ezh2 re-sensitized the LLC_4M cells to gemcitabine-induced cell death (Fig. 5H, I) and recovered the Gpx4 suppression in the LLC_4M cells (Fig. 5G). The database (GSE233468) analysis also showed that RIPK3 expression is higher upon EZH2 inhibitors treatment, including GSK126 and EPZ6438 (Fig. S4C). Enhanced EZH2 expression suppressed RIPK3 expression and rendered cells resistant to GEM treatment in HCC-827 cells (Fig. S4D, E). Therefore, our data collectively indicated that EZH2-mediated hypermethylation of the RIPK3 promoter occurred in lung cancer cells following long-term CS exposure.

**Fig. 5.**
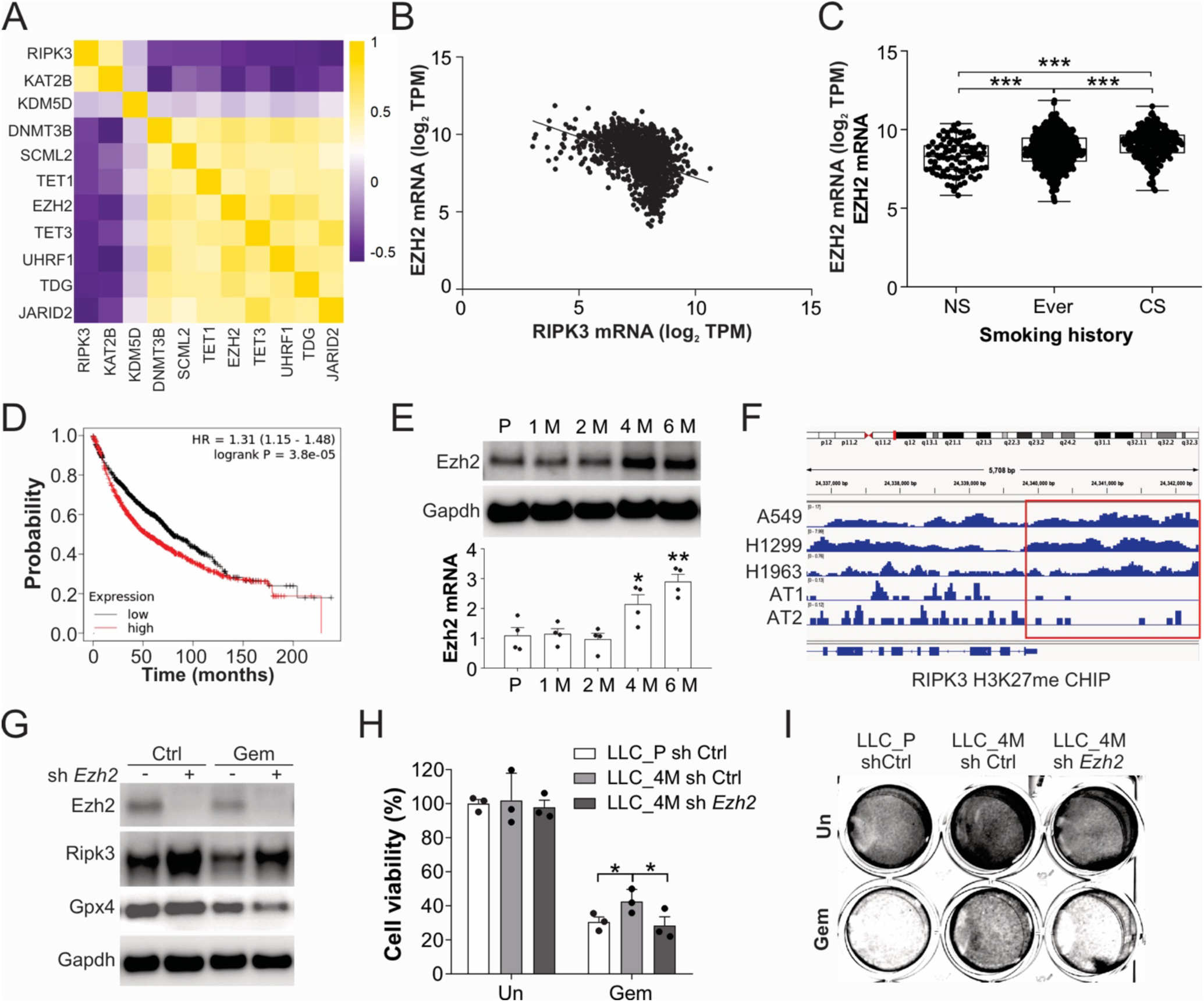
EZH2 suppresses RIPK3 expression upon long-term CS exposure. (A) The correlation of the TOP 10 epigenetic modification-related gene expression with RIPK3 expression was analyzed using the TCGA database. (B) The correlation of EZH2 and RIPK3 expression was analyzed using the TCGA database. Slope=-0.5263, R^2^=0.1710, p<0.001. (C) The mRNA level of EZH2 in lung cancer patients of never smokers (NS, n=93), ever smokers (n=636), and current smokers (CS, n=252). (D) Kaplan-Meier curves comparing overall survival (OS) in lung cancer patients with high or low EZH2 expression (https://kmplot.com/analysis/). (E) LLC cells were exposed to 4% CSE for 8 hours/day over 1, 2, 4, and 6 months. The expression of Ezh2 at the protein (upper) and mRNA (lower) levels was analyzed. (F) The binding of H3K27me3 to the RIPK3 promoter in different cells was analyzed. (G) LLC_4M cells stably transfected with sh Ctrl or sh Ezh2 shRNA were treated with 200 ng/ml Gem for 24h. The expression of the indicated proteins was analyzed by western blot. (H, I) The LLC_P and LLC_4M cells stably transfected with sh Ctrl or sh Ezh2 shRNA were treated with 200 ng/ml Gem for 24h. Cell viability was assessed by the MTT assay (H) and crystal violet staining (I). The experiments for (E, G-I) were repeated 3 times. Values in bar graphs are presented as means ± SD. *P* (t-test), *, p<0.05; **, p<0.01, ***, p<0.001.

### Targeting EZH2 overcomes the chemoresistance caused by Ripk3 downregulation

To explore the potential role of EZH2 targeting in overcoming CS-induced chemoresistance, we treated LLC_4M cells with the EZH2 inhibitor, GSK126. The pre-treatment with GSK126 enhanced the killing effect of gemcitabine on LLC_4M cells (Fig. 6A, B). A combination of GSK126 also rescued Ripk3 expression and resensitized it to Gpx4 suppression upon gemcitabine treatment (Fig. 6C). To further confirm the effect of targeting EZH2 *in vivo*, we implanted LLC_4M cells into the flanks of C57BL/6J mice and treated the mice with gemcitabine in combination with GSK126. While GSK126 alone didn’t significantly suppress the tumor growth of LLC_4M tumor, its combination with Gemcitabine significantly enhanced the tumor suppressive effect on LLC_4M cells (Fig. 6D). GSK126 treatment recovered the Ripk3 expression in the LLC_4M syngeneic tumors (Fig. 6E) and enhanced the gemcitabine-induced Gpx4 suppression (Fig. 6E, F). Given that ferroptosis has been reported as a form of immunogenic cell death[26], we further assess Cd8 immune cell accumulation in lung tumor tissues. We found that Gemcitabine treatment increased the Cd8 accumulation in the LLC_4M tumors (Fig. 6F, G). Therefore, our data indicate that targeting EZH2 can overcome chemoresistance induced by long-term CS exposure *in vitro* and *in vivo*.

**Fig. 6.**
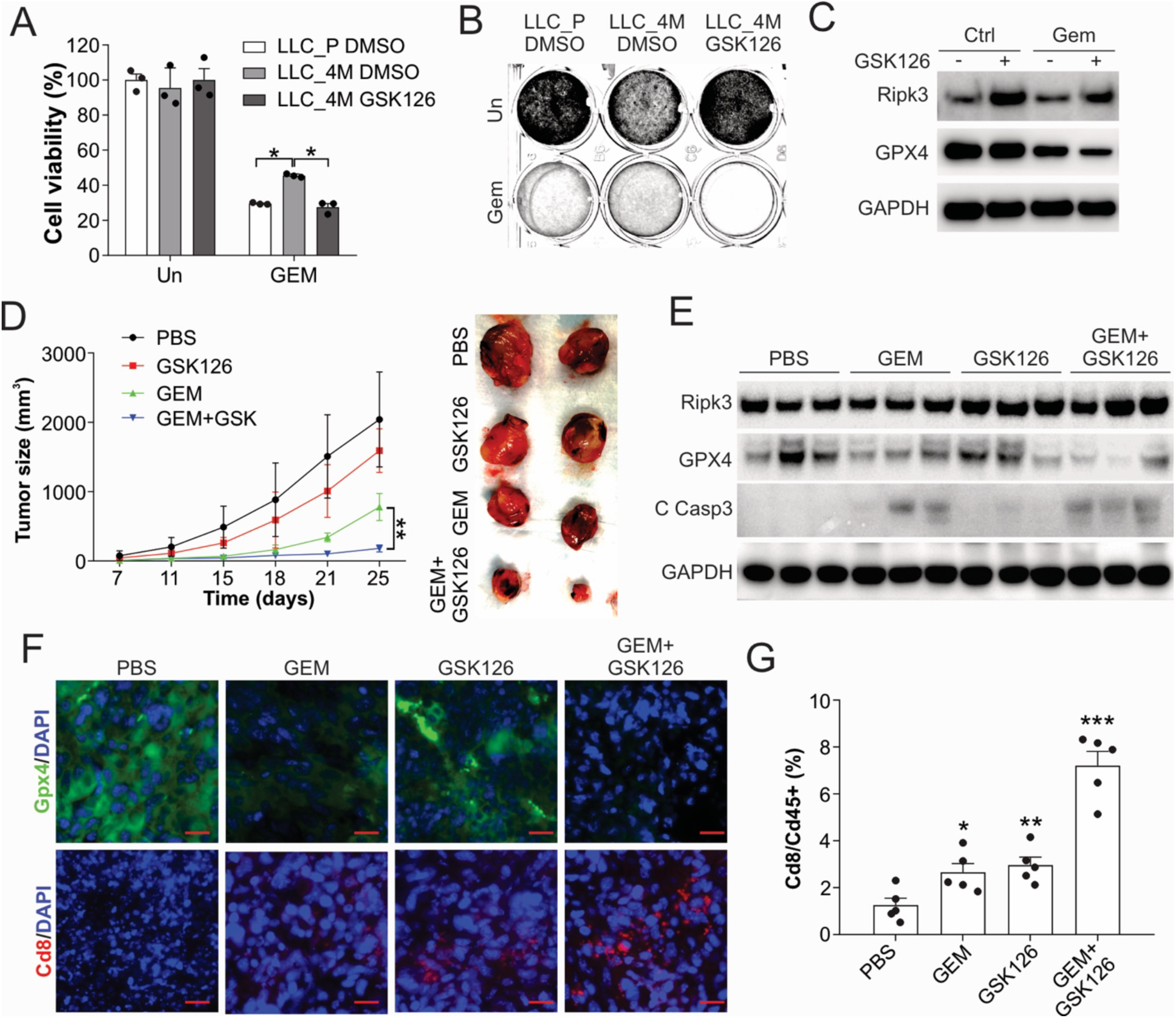
Targeting EZH2 overcomes the chemoresistance caused by CS exposure. (A, B) The LLC_P and LLC_4M cells were treated with 200 ng/ml Gem and/or 10 μM GSK126 for 24h. Cell viability was assessed by the MTT assay (A) and crystal violet staining (B). (C) LLC_4M cells were treated with 200 ng/ml Gem and/or 10 μM GSK126 for 24h. The expression of the indicated proteins was analyzed by western blot. (D-G) WT female C56B/6J mice were subcutaneously injected with LLC_4M cells (1X10^6^) (n=5 for each group). 7 days later, the mice were treated with Gem (25 mg/kg, 3 times/week) and/or GSK126 (50 mg/kg, 3 times/week). (D) The tumor size was measured on the indicated days. Left: growth curves were plotted; right: representative images of the tumors in each group. (E) The expression of the indicated proteins in each tumor group was analyzed by Western blot. (F) The representative image of Gpx4 and Cd8 immunofluorescence staining in each group of tumors. Scale bar, 25 μm. (G) Cd8 T cell accumulation in each tumor group was analyzed by flow cytometry. The experiments for (A-C) were repeated 3 times. Values in bar graphs are presented as means ± SD. *P* (t-test), *, p<0.05; **, p<0.01, ***, p<0.001.

## Discussion

Long-term CS exposure is a major contributor to chemoresistance in lung cancer treatment[27, 28]. The cytotoxic effects of most chemotherapeutic agents primarily rely on inducing apoptosis[3, 29]. However, cancer cells can escape this program, leading to uncontrolled proliferation, therapeutic resistance, and recurrence of cancer[5, 30]. Recent studies showed that other forms of programmed cell death, including necroptosis and ferroptosis, can be a target for cancer therapy to bypass apoptosis resistance[6–9]. Here, we show that long-term CS exposure suppresses expression of RIPK3, a key molecular modulator of necroptosis, in lung cancer cells, thereby impairing chemotherapy-induced cell ferroptosis instead of necroptosis. Mechanistically, we identify EZH2 as a key mediator of this effect. Chronic CS exposure upregulates EZH2, which promotes epigenetic repression of RIPK3 and is associated with the promoter H3K27 hypermethylation. Importantly, genetic or pharmacologic inhibition of EZH2 restores RIPK3 expression and resensitizes tumor cells to chemotherapy-induced cell death (Fig. 7). These findings establish an EZH2–RIPK3 axis as a critical pathway linking chemoresistance caused by smoking.

**Fig. 7.**
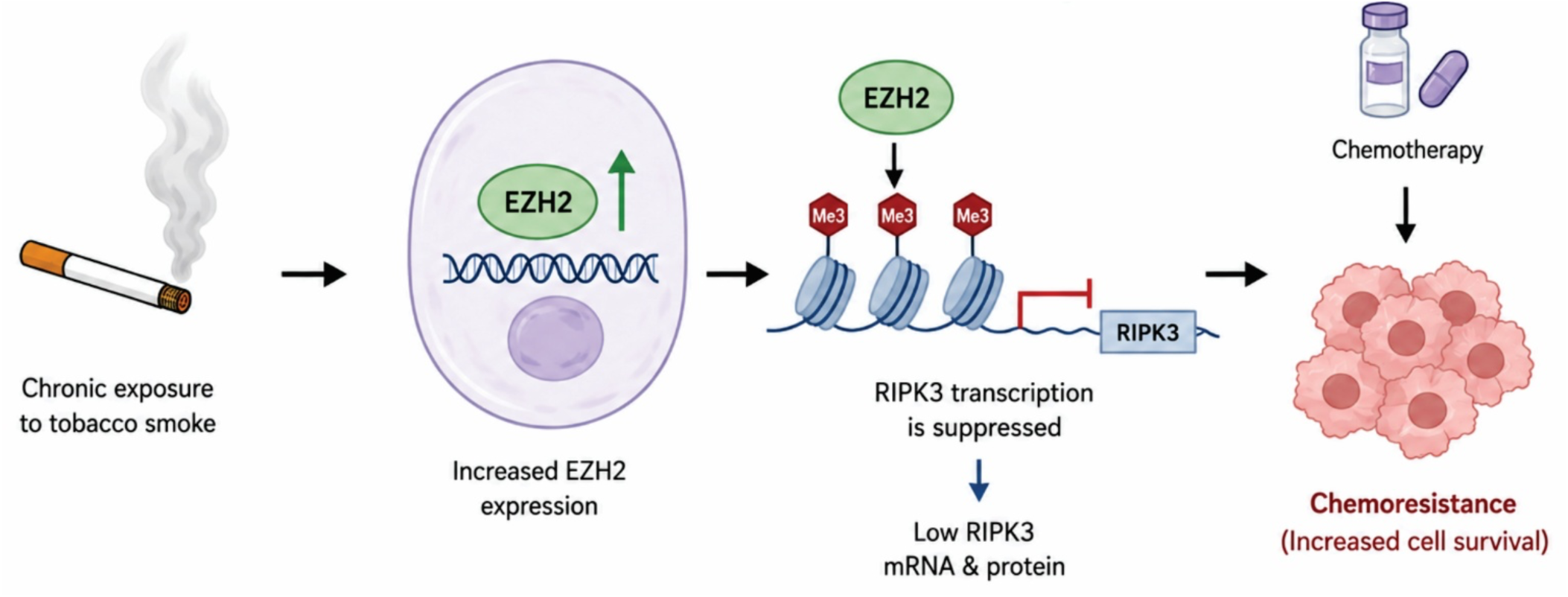
Model of Action.

Although RIPK3 silence has been observed in lung cancer[31], the underlying regulatory mechanisms remain incompletely understood, especially in those patients with a smoking history. In this study, we identified EZH2, a key epigenetic regulator frequently overexpressed in lung tumors[32], as a critical mediator of RIPK3 repression upon long-term CS exposure. EZH2 has emerged as an important therapeutic target in lung cancer, with multiple EZH2 inhibitors currently undergoing clinical evaluation[33]. Elevated EZH2 expression has been associated with smoking-induced lung cancer progression, including enhanced tumor proliferation and metastasis[34, 35]. However, its contribution to smoking-associated chemoresistance has remained largely unexplored. In the present study, we found that long-term CS exposure upregulates EZH2 expression and suppresses RIPK3 expression. Given that EZH2 catalyzes trimethylation of histone H3 lysine 27 (H3K27me3), a well-established repressive chromatin mark, we further demonstrated that H3K27me3 is enriched at the RIPK3 promoter in multiple lung cancer cell lines with low RIPK3 expression, suggesting that EZH2 represses RIPK3 transcription through epigenetic silencing. Importantly, genetic depletion or pharmacological inhibition of EZH2 restored RIPK3 expression and reversed CS-induced chemoresistance. These findings are consistent with the established oncogenic role of EZH2 in driving epigenetic reprogramming and tumor progression and support the use of EZH2 inhibitors as a therapeutic strategy for those lung cancer patients with a smoking history.

Interestingly, the silencing of RIPK3 by long-term CS exposure suppressed chemotherapy-induced ferroptosis rather than necroptosis. As two distinct modalities of programmed cell death, ferroptosis and necroptosis exhibit intricate crosstalk through shared initiators and intersecting signaling cascades[26]. The most prominent feature of ferroptosis is iron-dependent lipid peroxidation[26], whereas necroptosis also generates ROS[36]. RIPK3 has been reported to induce ROS by phosphorylating proteins, thereby promoting mitochondrial dysfunction, metabolic reprogramming, and increasing mitochondrial ROS (mtROS) production[36, 37]. In this study, our data supported the idea that RIPK3 expression contributes to ROS generation induced by chemotherapy drugs, thereby promoting ferroptosis. However, the underlying mechanism by which RIPK3 triggers chemotherapy-induced ROS may still require further investigation.

In summary, our findings reveal a novel mechanism by which long-term CS exposure promotes lung cancer progression and chemoresistance through EZH2-mediated epigenetic silencing of RIPK3, thereby inhibiting cell death. Targeting this pathway with EZH2 inhibitors or hypomethylating agents may restore sensitivity to chemotherapy drugs. This study provides a strong rationale for developing combination therapeutic strategies for lung cancer patients with a history of smoking.

## Conflict of Interest

None of the authors has any conflict of interest to disclose.

## Data access statement

The data was available upon request.

## Acknowledgment

We thank Dr. Jessie Huang (University of Southern California) for providing the pHAGE-EZH2 plasmid. This work was supported by the U.S. NIH grant (R56HL162749, D. Chen), TRDRP award (T35KT9504, D. Chen), ACS pilot grant (#IRG –22-144-60, D. Chen), and start-up funds from the Keck School of Medicine of USC.

During the preparation of this work, the author(s) used ChatGPT (OpenAI, 2026) to generate methods for Fig. 5A and the model of action for Fig. 7. After using this tool/service, the author(s) reviewed and edited the content as needed and take(s) full responsibility for the content of the publication.

## Author contributions

DC designed and monitored the research. RL, AZ, and DC performed the experiments. JY and DC performed the bioinformatic data analysis. RL and DC analyzed the data and wrote the paper. GX provided suggestions.

## Notes

### Competing Interest Statement

The authors have declared no competing interest.

## References

[1] C. Centers for Disease, Prevention, Cigarette smoking-attributable morbidity---United States, 2000, MMWR Morb Mortal Wkly Rep, 52 (2003) 842–844.

[2] G.M. Videtic, L.W. Stitt, A.R. Dar, W.I. Kocha, A.T. Tomiak, P.T. Truong, M.D. Vincent, E.W. Yu, Continued cigarette smoking by patients receiving concurrent chemoradiotherapy for limited-stage small-cell lung cancer is associated with decreased survival, J Clin Oncol, 21 (2003) 1544–1549.

[3] A. Eastman, Activation of programmed cell death by anticancer agents: cisplatin as a model system, Cancer Cells, 2 (1990) 275–280.

[4] W.L. Heusch, R. Maneckjee, Signalling pathways involved in nicotine regulation of apoptosis of human lung cancer cells, Carcinogenesis, 19 (1998) 551–556.

[5] D. Chen, J. Yu, L. Zhang, Necroptosis: an alternative cell death program defending against cancer, Biochim Biophys Acta, 1865 (2016) 228–236.

[6] Y. Gong, Z. Fan, G. Luo, C. Yang, Q. Huang, K. Fan, H. Cheng, K. Jin, Q. Ni, X. Yu, C. Liu, The role of necroptosis in cancer biology and therapy, Mol Cancer, 18 (2019) 100.

[7] D. Chen, K. Ermine, Y.J. Wang, X. Chen, X. Lu, P. Wang, D. Beer-Stolz, J. Yu, L. Zhang, PUMA/RIP3 Mediates Chemotherapy Response via Necroptosis and Local Immune Activation in Colorectal Cancer, Mol Cancer Ther, 23 (2024) 354–367.

[8] X. Lu, D. Chen, M. Wang, X. Song, K. Ermine, S. Hao, A. Jha, Y. Huang, Y. Kang, H. Qiu, H.J. Lenz, S. Li, Z. Jin, J. Yu, L. Zhang, Depletion of oxysterol-binding proteins by OSW-1 triggers RIP1/RIP3-independent necroptosis and sensitization to cancer immunotherapy, Cell Death Differ, 32 (2025) 2038–2052.

[9] K. Liu, J. Huang, J. Liu, D.J. Klionsky, R. Kang, D. Tang, Induction of autophagy-dependent ferroptosis to eliminate drug-tolerant human retinoblastoma cells, Cell Death Dis, 13 (2022) 521.

[10] J. Yan, P. Wan, S. Choksi, Z.G. Liu, Necroptosis and tumor progression, Trends Cancer, 8 (2022) 21–27.

[11] H. Yang, Y. Ma, G. Chen, H. Zhou, T. Yamazaki, C. Klein, F. Pietrocola, E. Vacchelli, S. Souquere, A. Sauvat, L. Zitvogel, O. Kepp, G. Kroemer, Contribution of RIP3 and MLKL to immunogenic cell death signaling in cancer chemotherapy, Oncoimmunology, 5 (2016) e1149673.

[12] F. Sun, L. Li, P. Yan, J. Zhou, S.D. Shapiro, G. Xiao, Z. Qu, Causative role of PDLIM2 epigenetic repression in lung cancer and therapeutic resistance, Nat Commun, 10 (2019) 5324.

[13] D. Chen, A.D. Gregory, X. Li, J. Wei, C.L. Burton, G. Gibson, S.J. Scott, C.M. St Croix, Y. Zhang, S.D. Shapiro, RIP3-dependent necroptosis contributes to the pathogenesis of chronic obstructive pulmonary disease, JCI Insight, 6 (2021).

[14] D. Chen, J. Tong, L. Yang, L. Wei, D.B. Stolz, J. Yu, J. Zhang, L. Zhang, PUMA amplifies necroptosis signaling by activating cytosolic DNA sensors, Proc Natl Acad Sci U S A, 115 (2018) 3930–3935.

[15] M.S. Cline, B. Craft, T. Swatloski, M. Goldman, S. Ma, D. Haussler, J. Zhu, Exploring TCGA Pan-Cancer data at the UCSC Cancer Genomics Browser, Sci Rep, 3 (2013) 2652.

[16] J. Goffin, C. Lacchetti, P.M. Ellis, Y.C. Ung, W.K. Evans, C. Lung Cancer Disease Site Group of Cancer Care Ontario’s Program in Evidence-Based, First-line systemic chemotherapy in the treatment of advanced non-small cell lung cancer: a systematic review, J Thorac Oncol, 5 (2010) 260–274.

[17] L. Sun, H. Wang, Z. Wang, S. He, S. Chen, D. Liao, L. Wang, J. Yan, W. Liu, X. Lei, X. Wang, Mixed lineage kinase domain-like protein mediates necrosis signaling downstream of RIP3 kinase, Cell, 148 (2012) 213–227.

[18] K. Lai, J. Wang, S. Lin, Z. Chen, G. Lin, K. Ye, Y. Yuan, Y. Lin, C.Q. Zhong, J. Wu, H. Ma, Y. Xu, Sensing of mitochondrial DNA by ZBP1 promotes RIPK3-mediated necroptosis and ferroptosis in response to diquat poisoning, Cell Death Differ, 31 (2024) 635–650.

[19] G.B. Koo, M.J. Morgan, D.G. Lee, W.J. Kim, J.H. Yoon, J.S. Koo, S.I. Kim, S.J. Kim, M.K. Son, S.S. Hong, J.M. Levy, D.A. Pollyea, C.T. Jordan, P. Yan, D. Frankhouser, D. Nicolet, K. Maharry, G. Marcucci, K.S. Choi, H. Cho, A. Thorburn, Y.S. Kim, Methylation-dependent loss of RIP3 expression in cancer represses programmed necrosis in response to chemotherapeutics, Cell Res, 25 (2015) 707–725.

[20] A.E. Teschendorff, Z. Yang, A. Wong, C.P. Pipinikas, Y. Jiao, A. Jones, S. Anjum, R. Hardy, H.B. Salvesen, C. Thirlwell, S.M. Janes, D. Kuh, M. Widschwendter, Correlation of Smoking-Associated DNA Methylation Changes in Buccal Cells With DNA Methylation Changes in Epithelial Cancer, JAMA Oncol, 1 (2015) 476–485.

[21] M.A. Honer, B.I. Ferman, Z.H. Gray, E.A. Bondarenko, J.R. Whetstine, Epigenetic modulators provide a path to understanding disease and therapeutic opportunity, Genes Dev, 38 (2024) 473–503.

[22] H. Ashoor, C. Louis-Brennetot, I. Janoueix-Lerosey, V.B. Bajic, V. Boeva, HMCan-diff: a method to detect changes in histone modifications in cells with different genetic characteristics, Nucleic Acids Res, 45 (2017) e58.

[23] Z. Zhao, A.P. Szczepanski, N. Tsuboyama, H. Abdala-Valencia, Y.A. Goo, B.D. Singer, E.T. Bartom, F. Yue, L. Wang, PAX9 Determines Epigenetic State Transition and Cell Fate in Cancer, Cancer Res, 81 (2021) 4696–4708.

[24] V.M. Fava, M. Dallmann-Sauer, M. Orlova, W. Correa-Macedo, R. Olivenstein, C.T. Costiniuk, J.P. Routy, L.B. Barreiro, E. Schurr, Continuous antiretroviral therapy induces progressive senescence-like reprogramming of alveolar macrophages, Front Immunol, 17 (2026) 1805936.

[25] Y. Tsutsui, A. Masui, S. Konishi, T. Tsujimura, M. Iwasaki, T. Yamamoto, S. Gotoh, Human iPSC-based Modeling of Pulmonary Fibrosis Reveals p300/CBP Inhibition Suppresses Alveolar Transitional Cell State, Nat Commun, 17 (2026) 1214.

[26] W. Gao, X. Wang, Y. Zhou, X. Wang, Y. Yu, Autophagy, ferroptosis, pyroptosis, and necroptosis in tumor immunotherapy, Signal Transduct Target Ther, 7 (2022) 196.

[27] J. Zhang, O. Kamdar, W. Le, G.D. Rosen, D. Upadhyay, Nicotine induces resistance to chemotherapy by modulating mitochondrial signaling in lung cancer, Am J Respir Cell Mol Biol, 40 (2009) 135–146.

[28] A.S. Tsao, D. Liu, J.J. Lee, M. Spitz, W.K. Hong, Smoking affects treatment outcome in patients with advanced nonsmall cell lung cancer, Cancer, 106 (2006) 2428–2436.

[29] D. Chen, L. Ming, F. Zou, Y. Peng, B. Van Houten, J. Yu, L. Zhang, TAp73 promotes cell survival upon genotoxic stress by inhibiting p53 activity, Oncotarget, 5 (2014) 8107–8122.

[30] R.W. Johnstone, A.A. Ruefli, S.W. Lowe, Apoptosis: a link between cancer genetics and chemotherapy, Cell, 108 (2002) 153–164.

[31] D. Agrawal, K. Cisarova, S. Vosberg, F. Allmendinger, E. Munkhbaatar, N. Dandachi, F.J. Fernandez Hernandez, M. Tonietto, V. Jager, M. Anton, E.C. Keller, M. Jesinghaus, A.L. Meinhardt, V. Haefner, T. Stoeger, K. Steiger, N. McGranahan, M.A. Dengler, A. Wahida, P.J. Jost, Aberrant methylation limits antitumoral inflammation in lung adenocarcinoma by restricting RIPK3 expression, Sci Adv, 12 (2026) eadz9227.

[32] F.F. Hu, H. Chen, Y. Duan, B. Lan, C.J. Liu, H. Hu, X. Dong, Q. Zhang, Y.M. Cheng, M. Liu, A.Y. Guo, C. Xuan, CBX2 and EZH2 cooperatively promote the growth and metastasis of lung adenocarcinoma, Mol Ther Nucleic Acids, 27 (2022) 670–684.

[33] N.J. Choudhury, W.V. Lai, A. Makhnin, G. Heller, J. Eng, B. Li, I. Preeshagul, F.C. Santini, M. Offin, K. Ng, P. Paik, C. Larsen, M.S. Ginsberg, Y. Lau, X. Zhang, M.K. Baine, N. Rekhtman, C.M. Rudin, A Phase I/II Study of Valemetostat (DS-3201b), an EZH1/2 Inhibitor, in Combination with Irinotecan in Patients with Recurrent Small-Cell Lung Cancer, Clin Cancer Res, 30 (2024) 3697–3703.

[34] H. Huang, C. Ding, W.H. Zhao, H.B. Zhang, Z.X. Zhao, X.G. Li, Y.J. Wang, P.J. Chen, B.S. Li, X.B. Li, Y.W. Li, H.Y. Liu, J. Chen, Nicotine promotes the progression and metastasis of non-small cell lung cancer by modulating the OTUB1-c-Myc-EZH2 axis, Acta Pharmacol Sin, 46 (2025) 2509–2521.

[35] K. Fan, B.H. Zhang, D. Han, Y.C. Sun, EZH2 as a prognostic-related biomarker in lung adenocarcinoma correlating with cell cycle and immune infiltrates, BMC Bioinformatics, 24 (2023) 149.

[36] Z. Yang, Y. Wang, Y. Zhang, X. He, C.Q. Zhong, H. Ni, X. Chen, Y. Liang, J. Wu, S. Zhao, D. Zhou, J. Han, RIP3 targets pyruvate dehydrogenase complex to increase aerobic respiration in TNF-induced necroptosis, Nat Cell Biol, 20 (2018) 186–197.

[37] Y. Zhou, Y. Xiang, S. Liu, C. Li, J. Dong, X. Kong, X. Ji, X. Cheng, L. Zhang, RIPK3 signaling and its role in regulated cell death and diseases, Cell Death Discov, 10 (2024) 200.

