## Supplemental Figures for "Suppression of RIPK3 by EZH2 contributes to cigarette smoking-induced chemoresistance in lung cancer"

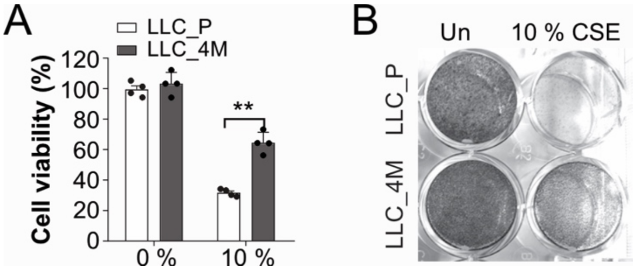


**Fig. S1. Long-term CS exposure leads to resistance to CS-induced cell death.**
The LLC_P and LLC_4M cells were treated with 10 % CSE for 24 h. The change in cell viability was analyzed using the MTT assay (A) and crystal violet staining (B). *P* (t-test), ****** <0.01.

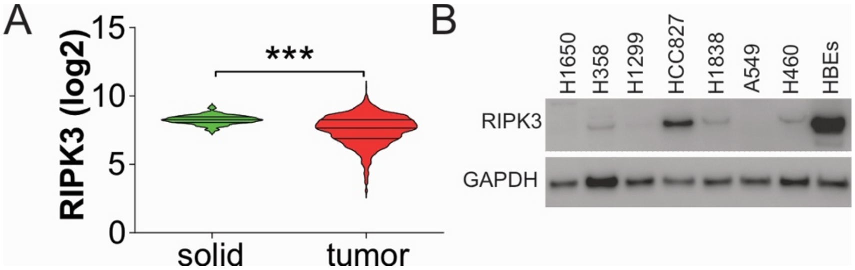


**Fig. S2. Lower RIPK3 expression in the lung tumors.**
(A) The mRNA level of RIPK3 in normal lung tissues and lung tumors from the TCGA database. (B) The western blot of RIPK3 in the indicated cell lines. *P* (t-test), *** <0.001.

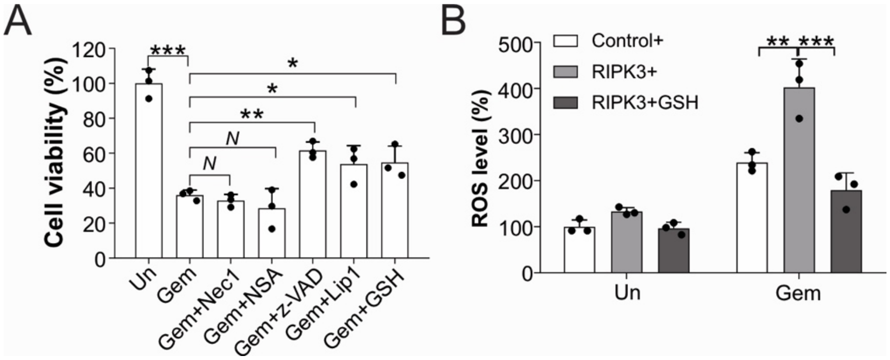


**Fig. S3. RIPK3 mediates Gem-induced ROS**

(A) The H460 were treated with 50 ug/ml Gem in combination with Nec-1 (2μM), NSA (2μM), z-VAD (10 μM), Lip1 (2 μM), and GSH (5 mM) for 24 h. Cell viability was analyzed by the MTT assay. (B) The LLC_4M cells transfected with pcDNA3.1 or pcDNA3.1 HA-Ripk3 were treated with 200 ng/ml Gem in combination with GSH (5 mM) for 24h. The ROS level in cells was measured using the H2DCFDA assay. Each experiment was repeated 3 times. Values in bar graphs are presented as means ± SD. N, $p>0.05$; *, $p<0.05$; **, $p<0.01$; ***, $p<0.001$.

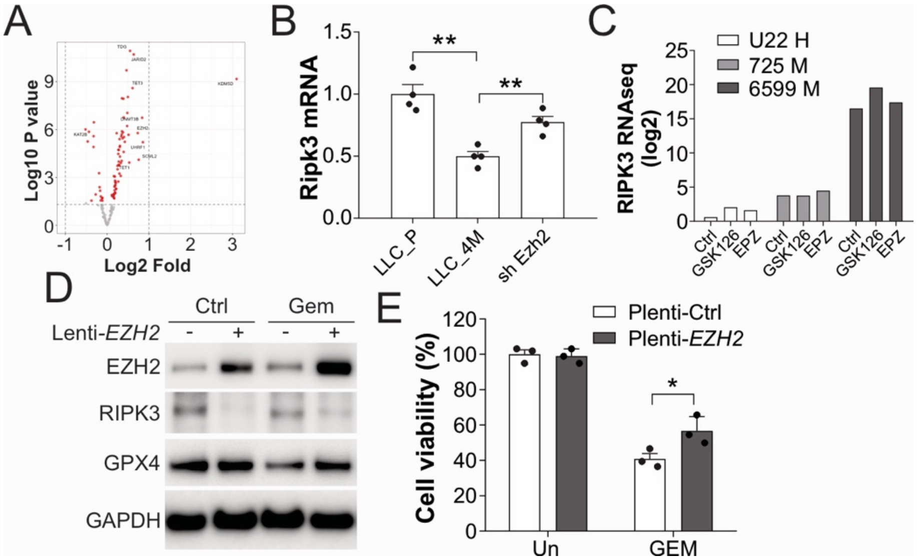


**Fig. S4. RIPK3 expression was modified in transcriptional level.**

(A) Differential expression of genes related to epigenetic modification between never smokers and current smokers in lung cancer patients based on the TCGA database. The top 10 differentially expressed genes were labeled. (B) The Ripk3 mRNA in LLC_P or LLC_4M cells stably transfected with sh Ctrl or sh *Ezh2* shRNA. (C) The database (GSE233468) analysis of RIPK3 mRNA changes in the lung tumor organoids treated with EZH2 inhibitors, GSK126, and EPZ6438. (C, D) The HCC827 cells transfected with a control or an EZH2-overexpressing lentivirus were treated with 50 ug/ml Gem for 24 h. (D) The expression of indicated proteins was analyzed by western blot. (E) The cell viability was analyzed by MTT assay. Each experiment was repeated 3 times. Values in bar graphs are presented as means ± SD. *p* (t-test), *, p<0.05; **, p <0.01.
